# Anti-*Candida* and antibiofilm activity of epinecidin-1 and its variants

**DOI:** 10.1101/2024.08.31.610647

**Authors:** Sivakumar Jeyarajan, Anbarasu Kumarasamy

## Abstract

To boost the stability and antimicrobial efficacy of the antimicrobial peptide (AMP) epinecidin-1, we previously engineered two variants — variant-1 and variant-2—by substituting alanine and histidine residues with lysine. This modification led to improved structural integrity and antibacterial function. Our current study builds on this foundation by assessing the anti-*Candida* capabilities of epinecidin-1 and its variants against test organisms *Candida albicans, Candida tropicalis*, and *Candida krusei*. Both variants exhibited potent anti-*Candida* activity, particularly in disrupting biofilm formation. Minimum Inhibitory Concentrations (MICs) were found to be decreased for both variants compared to the original epinecidin-1 peptide, with variant-2 exhibiting the strongest activity. Electron microscopy confirmed that the mechanism of action involves pore formation and the induction of reactive oxygen species in the *Candida* cell membrane. Computational analysis showed the peptides have a high tendency to interact with the *Candida* cell membrane proteins like Exo-B-(1,3)-Glucanase, Secreted aspartic proteinase (Sap) 1, and N-terminal domain adhesin: Als 9-2, to prevent biofilm development.

## 1. Introduction

Candidiasis, or *Candida* infections, have become an increasingly significant global health concern in the recent decades due to their heightened resistance to antifungal drugs such as fluconazole, the echinocandins, and polyenes [1, 2] Multi-drug resistance mechanism in *Candida sp*. is a major challenge to global health. The World Health Organization (WHO) has issued warnings about the potential ineffectiveness of many *Candida*cidal drugs in the future, placing immunocompromised patients, such as those with AIDS or cancer, or patients undergoing immune suppression for transplants or other medical procedures, at higher risk of severe infections. Hence there is a compulsive need for effective anti-*Candida* agents.

*Candida sp*. possess a defense-oriented outer membrane that consists of specialized lipids and sugars. They can alter this structure to evade the impact of drugs, which leads to resistance. Additionally, they go through a transformation process, often referred to as hyphal transformation, that assists them in circumventing the host’s immune system. These traits pose additional difficulties in orchestrating a successful treatment strategy for *Candida* infections. Despite advances in treating *Candida* infection, there is considerable interest in developing *Candida-*cidal agents with a new mode of action because of the biofilm formation, resistant development and non-responsiveness by invasive *Candida* species *Candida albicans, Candida tropicalis* and *Candida krusei [3]*. In recent times more studies are focused on studying cationic antimicrobial peptides (AMPs), which are toxic to *Candida* but not to normal mammalian cells. Hence AMPs are considered as potential alternatives to conventional *Candida*-cidals. Histatins[4], protonectin[5], cathelicidins from chicken and human namely CATH-2 and LL-37[6], N-terminal domain of bovine lactoferrin[7], protegrin etc. are some examples of potential AMPs having significant anti-*Candida* property. Currently more than 1400 AMPs with antifungal action in the APD3 database [8] .

Epinecidin-1, a cationic antimicrobial peptide (AMP) initially discovered in the grouper fish (*Epinephelus coioides*), displays a wide range of activity against bacteria, fungi, viruses, and protozoa [9-12] . As this peptide manifests such broad spectrum of activity, we have further augmented its structural and functional stability through the substitution of lysine at the alanine and histidine residues, as depicted in [11]. This modification resulted in variants with enhanced antibacterial performance. Given the negative charge and sugar composition analogous to the peptidoglycan of bacterial cell walls is also found in the fungal cell wall, it prompted us to explore the potential anti-*Candida* attributes of epinecidin-1 and its variants. In doing so, these variant peptides could possibly perform dual functions, serving as both antibacterial and anti-*Candida* agents. Our research findings reveal that these customized variants exhibit an increased *Candida*-cidal impact as like bactericidal.

## 2. Materials and Methods

### 2.1 Candida strains, peptides and Chemicals

To evaluate anti-*Candida* efficacy of the peptides, we conducted assays using *Candida albicans* (MTCC 227) along with biofilm-forming clinical isolates of *Candida tropicalis* (strain CA4) and *Candida krusei* (strain CA54) [13] as test organisms. These clinical isolates of *C. tropicalis* and *C. krusei* were generously provided by Dr. K. Natarajaseenivasan of the Department of Microbiology, Bharathidasan University, Tiruchirappalli, India.

The three peptides: epinecidin-1, and the computational designed lysine replaced ^*^variant-1, and variant-2, were purchased from (GenicBio Limited, Shanghai China), synthesized through the method of 9-fluorenylmethoxycarbonyl (Fmoc) solid-phase peptide synthesizer. All additional chemical reagents employed in the testing procedures were of analytical grade.

### 2.2 Variants of epinecidin-1

Epinecidin-1 and its lysine replaced variants were tested for anti-*Candida* property. Their sequences as reported in [11] with their corresponding site specific replaced amino acids are shown in Table 1.

**Table 1.** Sequence of epinecidin-1 and its variants. Lysine replaced residues are highlighted, bold, italicized, and underlined.

| Peptide name | Sequence |
| --- | --- |
| Epinecidin-1 | GFIFHIIKGLFHAGKMIHGLV |
| Variant 1- Replacement of H with K | GFIF<u>**K**</u>IIKGLF<u>**K**</u>AGKMI<u>**K**</u>GLV |
| Variant 2- Replacement of A with K | GFIFKIIKGLFK<u>**K**</u>GKMIKGLV |

### 2.3 Minimum inhibition concentration assay (MIC) and crystal violet staining

MIC was determined using the broth microdilution method as described by [3]. Briefly, *Candida* cells (1× 10^5^ CFU ml^-1^) were grown in Sabouraud dextrose broth (SDB) at 37° C for 24 h with and without peptides as control. Minimal inhibitory concentration (MIC) of the peptides was defined as the lowest concentration at which no visible turbidity was observed compared with control in which the *Candida* cells were not treated with peptides. The MICs assay was done in triplicates. After determining the MIC values, biofilm disruption activity was assayed by crystal violet staining as described earlier [3]. To measure the ability of the peptide to inhibit the formation of biofilm, 1× 10^5^ CFU ml^-1^ of *Candida* cells were seeded into 24-well titer plates with 1 ml of fresh SDB and the peptides at their respective MICs incubated overnight. The well containing culture and fresh media without peptide served as control. A microscope cover slip was inserted in each well, which served as a substratum for microbial attachment. After incubation, the media containing planktonic cells were aspirated from each well and 1 ml of 0.1 % (w/v) crystal violet was added and stained for 30 min followed by ethanol solubilization. The stained cover slips were examined under light microscope (×40).

### 2.4 Scanning Electron Microscopy (SEM)

To view the morphological changes of *Candida albicans* cells (MTCC 227) after treatment with epinecidin-1 and its variants, SEM was employed [14]. In brief, *Candida albicans* cells were grown to logarithmic phase with an inoculum size of (1× 10^5^ CFU ml^-1^) on a microscope cover slip which was inserted to each well of a 24 well dish. The *Candida albicans* were incubated with epinecidin-1, variant-1 and variant-2 at their MIC concentration of 128 μg ml^-1^, 64 μg ml^-1^ and 32 μg ml^-1^ along with Sabouraud dextrose broth medium as negative control. After incubation for 6 h at 37 °C (sub-lethal time point), the cells were washed twice with phosphate buffered saline (PBS) pH 7.4, and metabolically fixed with an equal volume of 5% (v/v) glutaraldehyde at 4°C overnight. The metabolically fixed cells were dehydrated with serial gradient of ethanol wash from 70% to 100%. The coverslips were then sputter coated and examined under a scanning electron microscope (VEGA3 TESCAN, Czech Republic).

### 2.5 Molecular docking study

The Protein Data Bank (PDB) files of *Candida* exo-beta-(1, 3)-glucanase (1CZ1), Sap1 (2QZW) and N-terminal domain of Als 9-2 (2Y7L) were obtained from the Research Collaboratory for Structural Bioinformatics (RSCB) protein data bank. The tertiary structure of the epinecidin-1 and designed variant peptides were obtained using i-TASSER [15]. The structures with minimum energy were selected for docking interaction using patchdock [16]. The patch dock output files were loaded into BIOVIA Discovery Studio (Dassault Systèmes, San Diego, 2021) for representation of the amino acids interacting with the *Candida* cell wall proteins and antimicrobial peptides. The amino acids interacting within 3□ distance are shown.

### 2.6 Measurement of cellular ROS production

Endogenous amounts of ROS were measured by fluorometric assay with 2′,7′-dichlorofluorescin diacetate (DCFHDA) as described in [5]. Briefly, *Candida* cells (1× 10^5^ CFU ml^-1^) were treated with or without peptides at their respective MIC for 24 h. After 24 h, the cells were incubated with 10 μM of DCFHDA for 1 h and washed with PBS pH 7.4. They were then visualized in fluorescent microscope (Accu-Scope, EXI-310, Commack, NY, USA) at 10x magnification and documented with green channel fluorescence intensities (excitation 488 nm and emission 525 nm respectively).

## 3. Results

### 3.1 Candidacidal activity of epinecidin-1 and its variants

The *Candida*-cidal activity of epinecidin-1 and its variants were studied using the broth dilution method. The Minimum Inhibitory Concentration (MIC) values obtained from the tests are shown in Table 2. For *Candida albicans*, the MIC values for epinecidin-1, variant-1 and variant-2 are 128 μg ml^-1^, 64 μg ml^-1^ and 32 μg ml^-1^, respectively. Therefore, variant-2 appears to exhibit the most effective anti-fungal properties, requiring the minimal concentration to inhibit the growth as shown in Figure 1.

**Table 2.** Minimal Inhibitory Concentration (MIC) of the peptides against tested *Candida* strains.

| Name of the organisms | Epinecidin-1 MIC (μg ml<sup>-1</sup>) | Variant-1 MIC (μg ml<sup>-1</sup>) | Variant-2 MIC (μg ml<sup>-1</sup>) |
| --- | --- | --- | --- |
| *C. albicans* | 128 | 64 | 32 |
| *C. tropicalis* | 256 | 64 | 32 |
| *C. krusei* | 128 | 128 | 64 |

**Figure 1.**
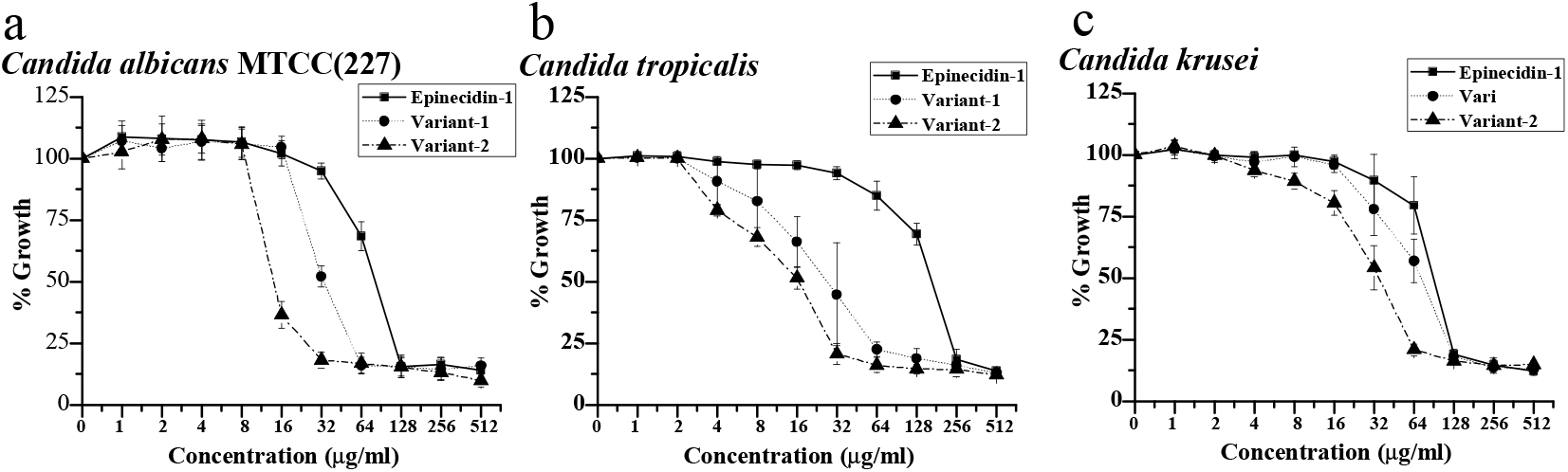
(Epinecidin-1, variant-1 and variant-2) action against a) *Candida albicans* (MTCC 227). More than 50% inhibition of growth was observed at 128 μg ml^-1^, 64 μg ml^-1^ and 32 μg ml^-1^. b) *Candida tropicalis;* epinecidin-1, variant-1 and variant-2 showed MIC of 256 μg ml^-1^, 64 μg ml^-1^, and 32 μg ml^-1^ respectively. c) *Candida krusei*; epinecidin-1 and variant-1 showed MIC of 128 μg ml^-1^ and variant-2 showed MIC of 64 μg ml^-1^. Cell density was measured at 595 nm after 24 h. Peptide concentration of 0 μg ml^-1^ denotes the control experiment.

*Candida tropicalis* showed similar patterns of sensitivity, with the respective MIC values for epinecidin-1, variant-1 and variant-2 (256 μg ml^-1^, 64 μg ml^-1^ and 32 μg ml^-1^). The trend continues with *Candida krusei*. The MIC values of epinecidin-1 and variant-1 against *Candida krusei* are 128 µg ml^-1^ and for variant-2 is 64 µg ml^-1^ respectively.

After determining the MIC, the ability of the peptides to disrupt the biofilm formed on glass cover slips was determined by crystal violet staining. *Candida* cells (*Candida albicans, Candida tropicalis*, and *Candida krusei*) were cultured with and without the peptides (at their respective MIC values) for a period of 24 h. After the treatment period, the coverslips containing the *Candida* cells were stained with crystal violet, and then examined under an inverted 40x light microscopy.

The results, as shown in Figure 2, provide visible evidence of the peptide’s impact on biofilm formation. Control samples (*Candida* cells without peptides) showed dense cell clusters indicative of substantial biofilm production. Conversely, peptide-treated *Candida* cells demonstrated major disruptions in the biofilm matrix and a significant reduction in cell numbers. This reduction suggests an inhibited biofilm formation capacity when the *Candida* cells were treated with these peptides. These results support the potential for epinecidin-1 and its variants to disrupt the formation of biofilms associated with *Candida* species.

**Figure 2.**
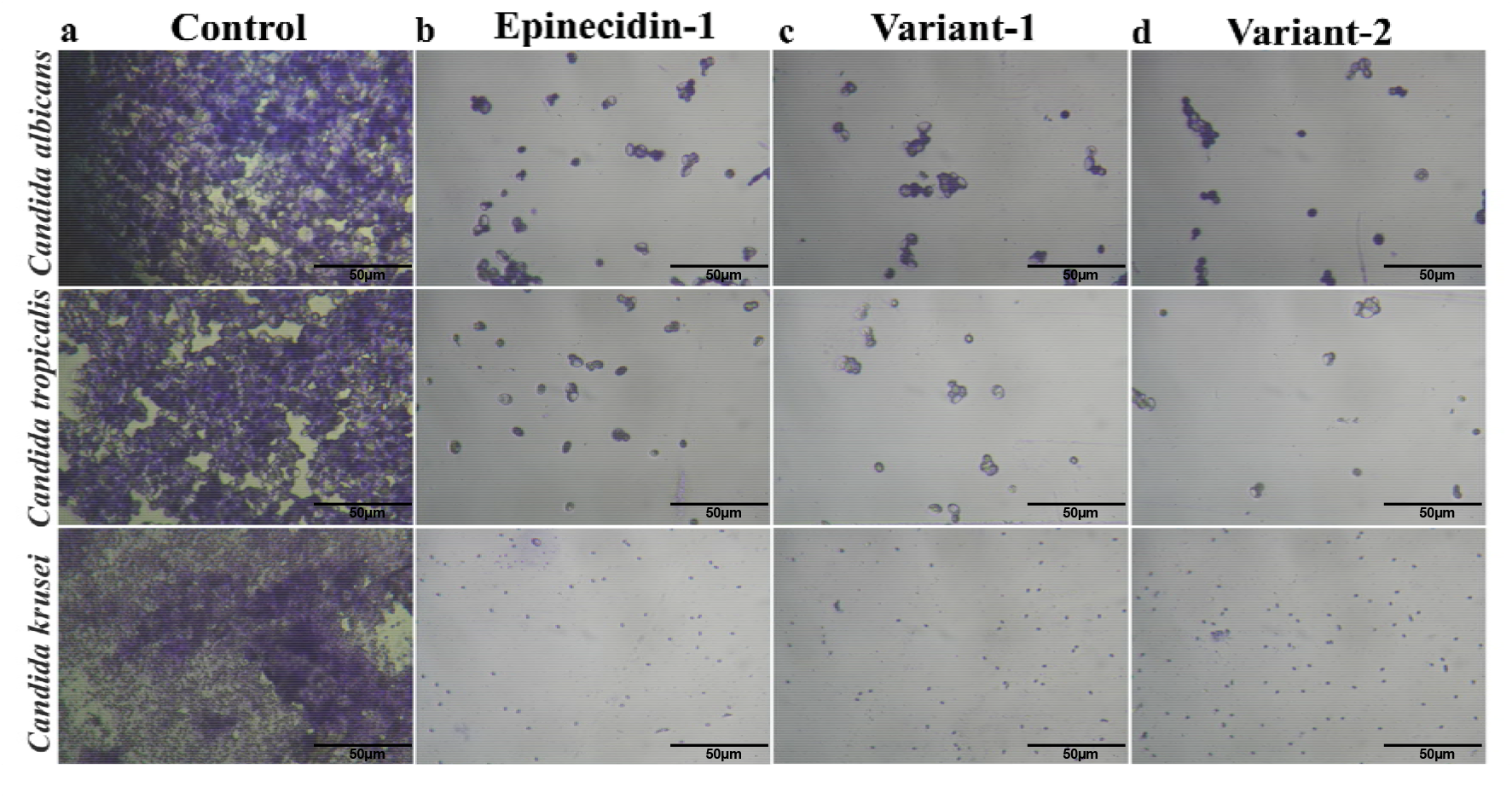
Light microscopic image of crystal violet stained a) *Candida albicans* (MTCC 227) and b) *C. tropicalis* and c) *C. Krusei* at 40x magnification. The cells were incubated with peptides at their respective MIC and incubated for 24 h. The cell number has decreased invariably for the *Candida* cells treated with peptides.

### 3.2 Ultrastructural morphology of Candida albicans membranes

Scanning Electron Microscopy (SEM) was utilized to investigate the impacts of epinecidin-1 and its variants on membrane penetration efficiency in the *C. albicans* cells as illustrated in Figure 3. Under SEM, the untreated control *C. albicans* cells exhibited typical features: oval shapes, smooth surfaces, polar buds, and bud scars. For examining the effects of the peptides on the *C. albicans* membrane, cells were incubated with 1x MIC of the peptides for 6 h, a sub-lethal time chosen due to the observation that cells typically detach after a 24 h treatment.

**Figure 3.**
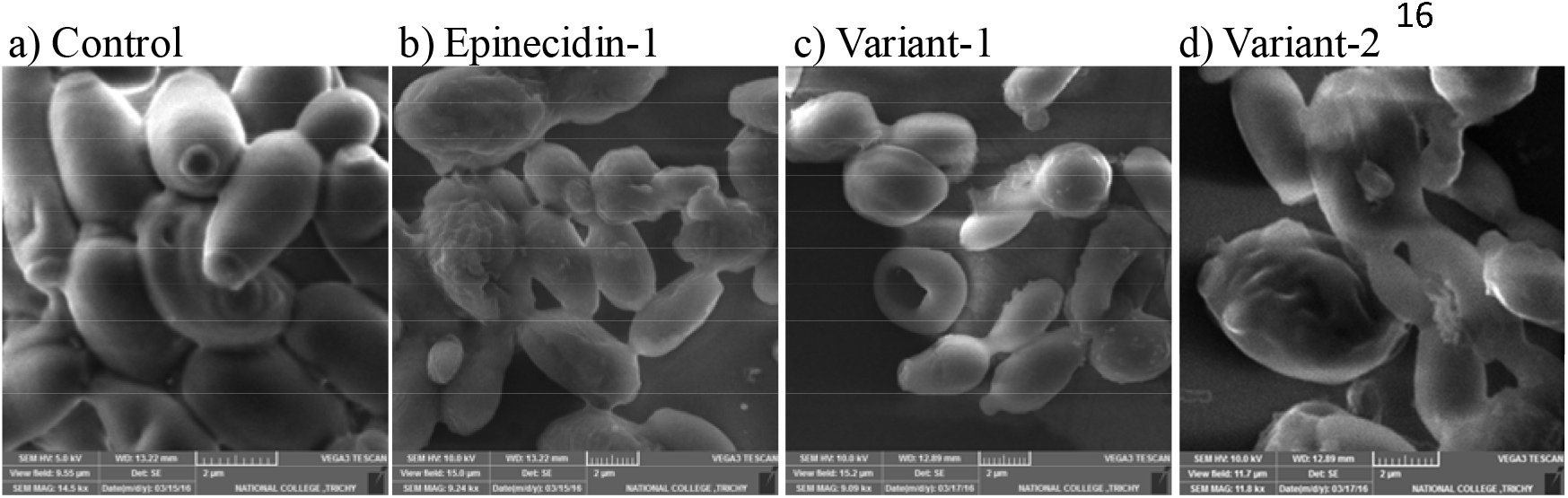
Scanning electron microscopic (SEM) image of *Candida albicans* (MTCC 227) (a) Control, and (b-d) peptide treated cells. The cells were treated with the peptides at their respective MIC and incubated for 6 h. The reduction in the number of cells noted as empty spaces between the cells and the disruption of cell membrane demonstrates the membrane disruption activity of the peptides. The peptide treated cells show aberrant surface with internally collapsed morphology due to cytoplasm leakage and appears.

The SEM images evidence that the peptide treatment altered the external cell morphology of *C. albicans*. When treated with epinecidin-1, the cells no longer appeared as smooth as the untreated cells; the membrane seems to inflate with a rough, uneven surface. Additionally, numerous vesicular structures were observed surrounding the treated cells. For the variant-1, the *Candida* cells displayed signs of shrinkage, potentially due to a loss of cytosolic content. Substantial perforations were apparent, suggesting the peptide-induced damage permeates the membrane, and more rounded cells were observed compared to the elongated cell morphology in the control. When treated with variant-2, the *Candida* cells endured severe damage, leading to the collapse of the cell wall and a noticeably bloated appearance, suggesting the possibility of a large pit on the opposite side. These findings show strong evidence of the antimicrobial activity of epinecidin-1 and its variants against *Candida*, resulting in significant membrane penetration potential and destabilization of the cell membrane.

### 3.3 Molecular interaction of peptides with *Candida* membrane proteins

To determine the capability of antimicrobial peptides to interact with the biofilm-forming proteins in the cell walls of *Candida*, computational methods such as Patch Dock and BIOVIA were used. The interface area of the peptide binding to three specific *Candida* cell wall proteins: a) Exo-B-(1,3)-Glucanase, b) Secreted Aspartic Proteinase (Sap) 1, and c) N-terminal domain of *Candida albicans* adhesin: Als 9-2 (Agglutinin Like Sequence Super Family 9; allelic isoform 2) were calculated and portrayed in Table 3. Data from PatchDock suggested that all the peptides bound to all the three selected target *Candida* cell wall proteins within a 3□ space and are represented in Figure 4. The amino acids of the *Candida* cell wall proteins interacting with the peptides are shown in white in Figure 4 and enumerated in Supplementary Table 1.

**Table 3.**
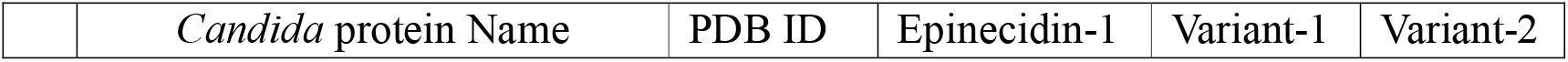
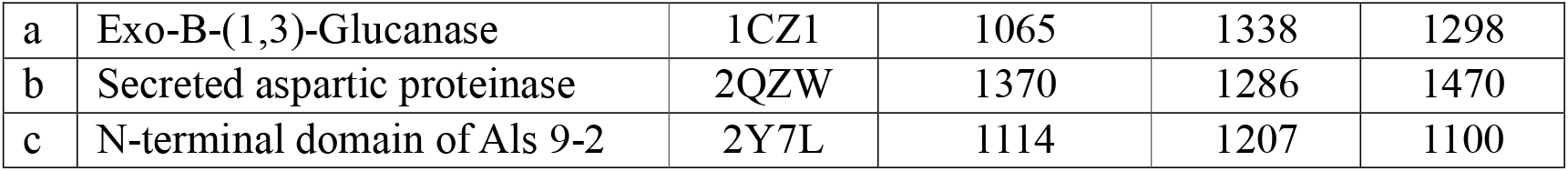
Interface area score of the peptides binding to selected *Candida* membrane proteins a) Exo-B-(1,3)-Glucanase, b) Secreted aspartic proteinase (Sap) 1 and c) N-terminal domain of Als 9-2 responsible for biofilm formation.

**Figure 4.**
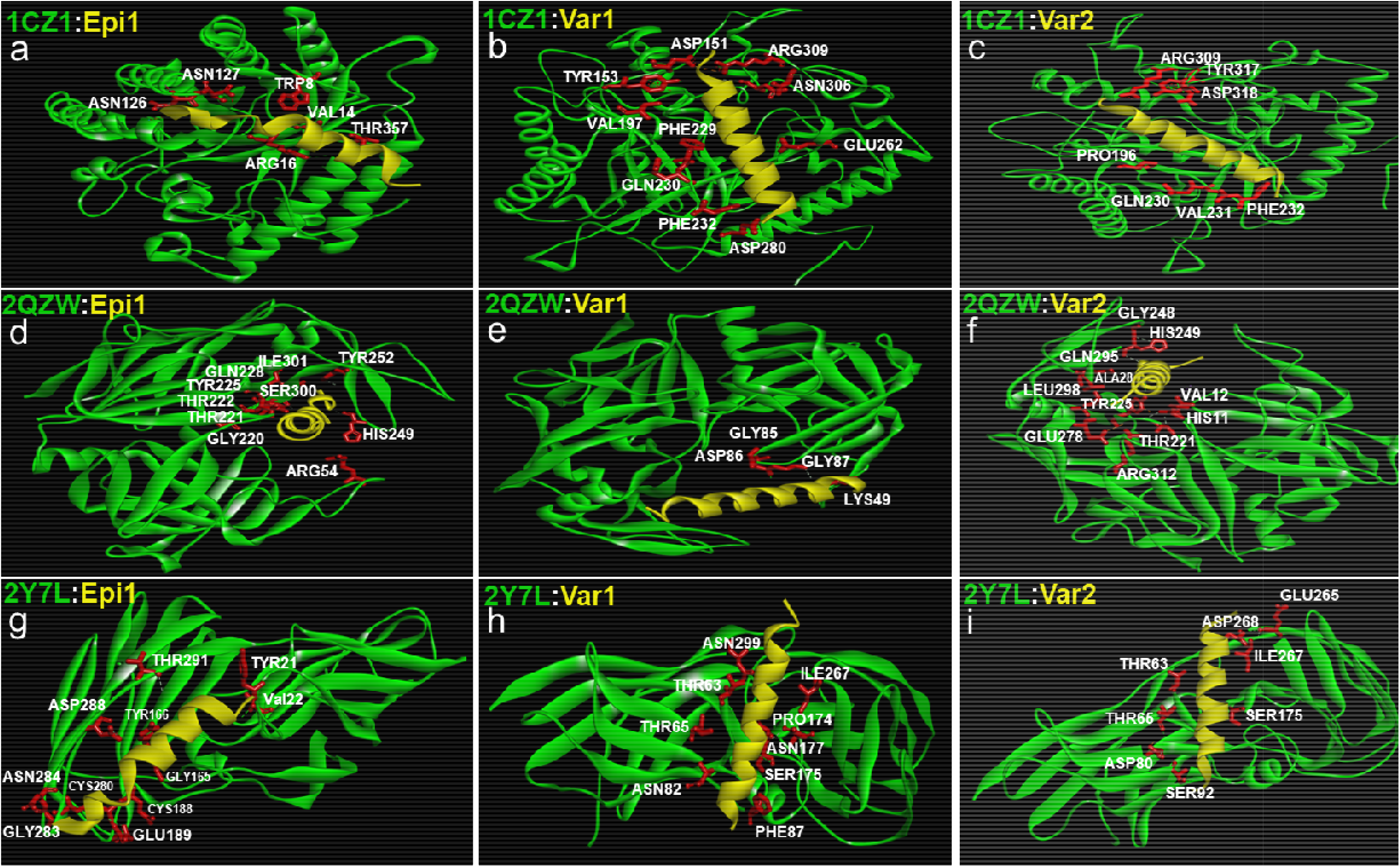
Docking interactions of epinecidin-1 and its variants with *Candida* Exo-B-(1,3)-Glucanase: PDB ID 1CZ1 (a-c), Secreted aspartic proteinase (Sap) 1: PDB ID 2QZW (d-f) and N-terminal domain of Als 9-2: PDB ID 2Y7L (g-i) biofilm forming membrane receptors. The *Candida* cell wall proteins are represented green in colour and the peptides are yellow in colour. The amino acids originating from the *Candida* cell wall proteins interacting with the peptides are represented in white with their three-letter code and their corresponding position. The amino acid side chain of the *Candida* protein (TYR, VAL, GLY, TYR, CYS, GLU, CYS, GLY, ASN, ASP, THR) acting as binding pocket for the peptides are shown red in colour.

A larger interface area signifies more interaction of the *Candida* cell wall proteins with the peptides. For Exo-B-(1,3)-Glucanase, variant-1 covers a larger interface area followed by variant-2 and epinecidin-1. The high positive charge of variant-1 and variant-2 leads to them having more interaction with the negatively charged amino acids of Exo-B-(1,3)-Glucanase. Variant-1 also interacts with hydrophobic and aromatic amino acids. Through Van Der Waals forces, Epinecidin-1 can interact with positively charged amino acids, while also binding with aromatic and hydrophobic amino acids like valine, proline, phenylalanine and glutamine.

In terms of interaction with secreted aspartic proteinase, variant-2 exhibits a greater interface area, followed by epinecidin-1 and variant-1. All three peptides interact with unique negatively charged amino acids and demonstrate interactions with hydrophobic and aromatic amino acids.

Examining the *Candida* adhesin protein namely (N-terminal domain of Als 9-2), variant-1 presents a higher interface surface area, while epinecidin-1 and variant-2 show similar values of interface area. Both variant-1 and variant-2 share mutual interactions with threonine 63, threonine 65, serine 175, and isoleucine 267 via hydrophobic and charge-based interactions. Most of the interactions for variant-2 occur through charge-based interactions. Meanwhile, in addition to these common interactions, variant-1 uniquely engages with asparagine using Van Der Waals forces. Epinecidin-1 exhibits a higher number of binding contacts with hydrophobic amino acids and fewer with both negatively and positively charged amino acids. These interactions primarily involve oppositely charged electrostatic interactions and likely charged Van Der Waals interactions. The binding site is also different from variants 1 and 2.

### 3.4 Epinecidin-1 and its variants induced ROS production

To determine whether the peptides have the ability to induce production of reactive oxygen species (ROS) production in *Candida* sp. we used dichlorofluorescein diacetate (DCFDA) after treatment with epinecidin-1 and its variants. DCFDA is a cell-permeant dye that is oxidized to yield fluorescence when exposed to ROS. The fluorescence can be monitored with the excitation wavelength of 488 nm and the emission wavelength of 525 nm. In Figure 5, the control cells (no treatment) showed a weak DCFDA fluorescence. The cells treated with the peptides for 24 h showed strong fluorescence intensity. Epinecidin-1 and variant-2 showed increased ROS production whereas variant-1 showed slightly less ROS production compared to epinecidin-1 and variant-2. These results strongly indicate that the peptides induce ROS production in *Candida* cells.

**Figure 5.**
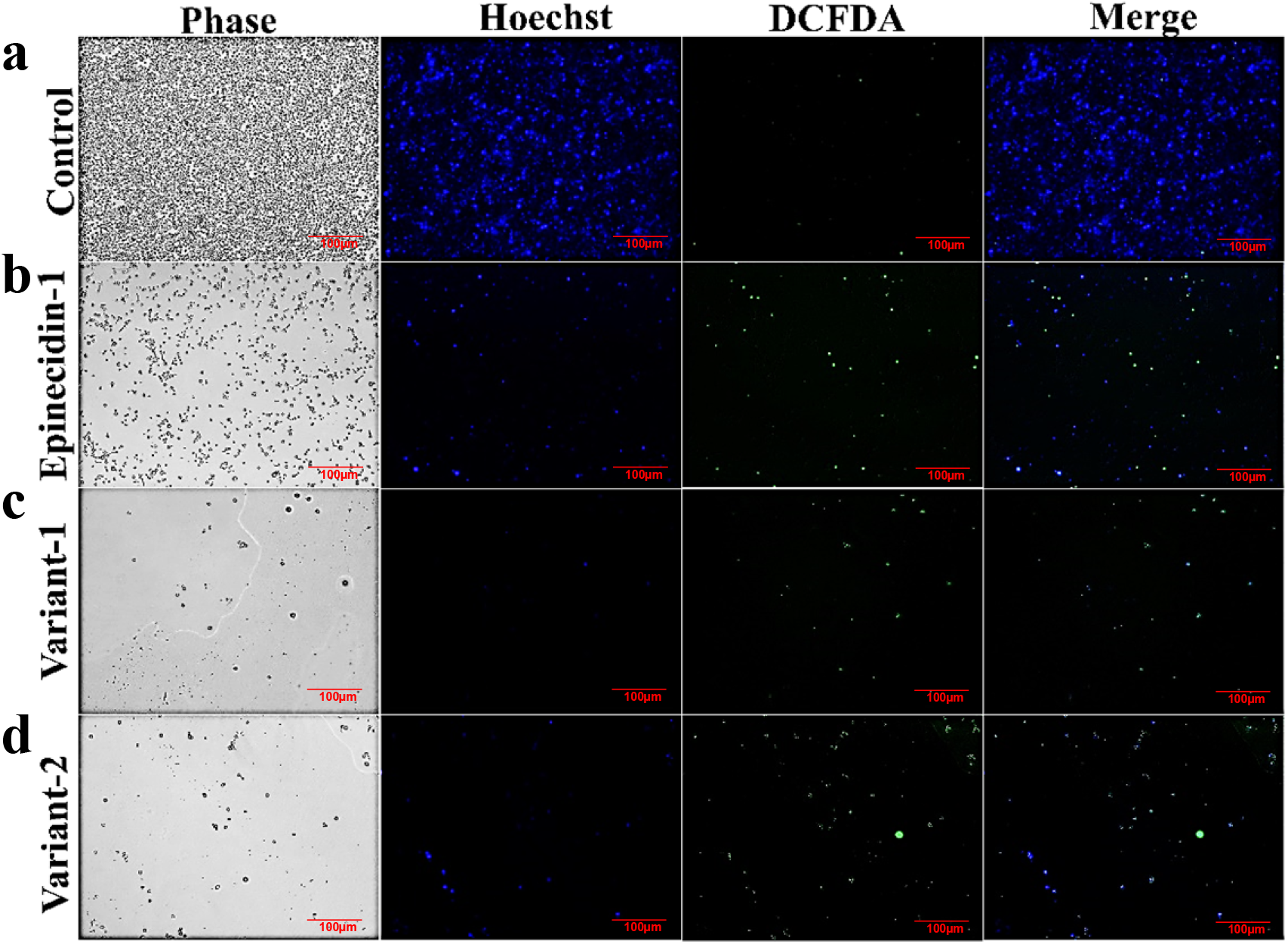
Effect of epinecidin-1 and its variants on the production of ROS by *Candida* cells (a) control cells, (b) epinecidin-1, (c) variant-1 and (d) variant-2 peptide treated cells. Logarithmically growing *C. albicans* cells (1×10^5^) were pre-incubated with 20 μM DCFHDA and grown for 24 h at 37°C. Cells were treated with and without peptides. After 24 h, the culture dish was washed with phosphate buffered saline pH 7.4 and observed under a fluorescent microscope. Hoechst staining was done alongside to determine the number of live cells. The images were captured at excitation wavelength of 488 nm and the emission wavelength of 525 nm.

## 4. Discussion

Candidiasis is a common and serious infection especially in the immune compromised patients and in the United States alone, 25,000 cases are reported annually and ranking as the fourth most frequent hospital-acquired infection [17] [18]. These infections often result in extended hospital stays, higher medical costs, and negatively affecting quality of life in patients. *Candida*, typically a harmless commensal on the skin and mucosal surfaces [19], can become a harmful pathogen in immunocompromised individuals, such as those with autoimmune diseases, diabetes, tuberculosis, AIDS, or those undergoing treatments like steroid administration for organ transplantation [20]. The risk is especially high for postoperative ICU patients [21], where outbreaks are more frequent and severe due to weakened immune systems which serves as conducive environment for *Candida* growth. Despite the use of antibiotics like Azoles, Echinocandins, and Polyenes, treatments are challenging due to the development of drug resistance [18].

During the COVID-19 pandemic, *Candida* infections were a leading cause of mortality, often affecting the lungs, throat, or bloodstream of COVID-19 patients due to their compromised immune systems, and potentially leading to sepsis [22, 23]. Conventional anti-*Candida* treatments were unsuitable for these secondary infections because their adverse effects in this context were unknown, highlighting the need for potent and natural antifungal treatments with lower risks of immune reactions.

*Candida albicans, Candida tropicalis*, and *Candida krusei* are tested due to their association with 95% of healthcare-associated invasive candidiasis worldwide, with substantial morbidity and mortality [24]. *C. albicans* is responsible for over 50% of *Candida* diseases and a leading cause of hospital-acquired bloodstream infections, with a mortality rate between 30%-60% [25]. *C. tropicalis*, linked to 40-60% of deaths from nosocomial infections, often develops resistance to azole antifungal agents [26]. *C. krusei*, although less prevalent, poses a high risk to patients with hematological malignancies and bone marrow transplants, and shows high resistance to fluconazole and amphotericin B, with significant morbidity, mortality and biofilm forming capabilities [27].

Antimicrobial peptides (AMPs) are short peptides; some of which exhibit helical or beta-sheet structures. These structural properties impart specific characteristics such as the ability to penetrate cells [11] [28, 29] and kill pathogens by forming aggregates [30-34]. The effectiveness of AMPs is influenced by factors such as net charge, hydrophobicity, and amphipathicity, with the amino acid composition being a critical determinant. Additionally, the helical structures of AMPs play a vital role in penetrating the fungal membrane, as emphasized by Sonesson *et al*. (2007)[35]. *Candida sp*. rely on their cell walls, comprised of components such as β-glucan, chitin, and phosphomannoproteins, for protection against environmental stress and antifungal drugs. These components have a negative charge, which makes them potential binding sites for positively charged AMPs. This interaction has been demonstrated in studies like that of Harris, M. *et al*. (2009) [36], where glycosylation knockouts of *C. albicans* resulted in a reduced negative charge and lower susceptibility to AMPs. It was observed that the binding of certain AMPs to the cell wall is crucial for their antifungal activity and here the charge and hydrophobicity are key factors in their function. Strategic design, guided by sequence databases, can be employed to engineer AMPs with enhanced specificity against microbes and reduced hemolytic activity. These tailored AMPs may be more appropriate for clinical applications. Within this realm, two variants of epinecidin-1, featuring lysine substitutions, have been developed, demonstrating increased lytic activity against *Candida sp*.

Epinecidin-1 inhibited the growth of *C. albicans* and *C. krusei* at 128 μg/mL, and *C. tropicalis* at a minimum inhibitory concentration (MIC) of 256 μg/mL. The histidine-to-lysine substituted variant, variant-1, inhibited all three *Candida* strains at 64 μg/mL, demonstrating a two-fold increase in activity against *C. albicans* and *C. krusei*, and a four-fold increase for *C. tropicalis*. The improved efficacy may be attributed to an increased helical propensity and a higher charge relative to epinecidin-1. These findings align with prior research on modified peptides, such as C3a peptides [35], the N-terminal domain of bovine lactoferrin[7], uperin 3.6 peptide[37], synthetic helical peptides [32], and KABT-AMP[3], which underscore similar structural features enhancing antimicrobial activity.

The variant-2 generated by replacement of weak hydrophobic alanine at 13^th^ position of variant-1 with charged amino acid lysine, inhibited the growth of *C. albicans* and *C. tropicalis* at 32 μg ml^-1^ and *C. krusei* at 64 μg ml^-1^. For *C. albicans*, there was a four-fold enhanced activity compared to epinecidin-1 and two-fold enhanced activity compared to variant-1. For *C. tropicalis* there was an eight-fold enhanced activity compared to epinecidin-1 and two-fold enhanced activity compared to variant-1. For *C. krusei*, there was two-fold enhancements compared to epinecidin-1 and similar for variant-1. The culminated enhanced activity of variant-2 would be from a large patch of positive charge formed by local electrostatic interactions [3, 9, 11] that mediate strong binding to cell membranes required for anti-*Candida* activity.

The majority of the pathogenic *Candida* species are protected by the formation of biofilms. During biofilm formation, an extracellular matrix is formed, encasing the microbial cell that prevents the antimicrobial substances reaching the cells [39]. Biofilms contribute significantly to microbe’s resistance against anti-fungal agents; disrupting them could enhance treatment effectiveness. Cells associated in biofilm formation requires 1000-fold greater concentration of fluconazole and amphotericin B to inhibit *Candida* biofilm than the planktonically grown strains [38, 39]. This feature plays a major role for persistent re-occurrence of infections and antibiotic resistance. Strikingly, for the *in vitro* peptide treated *Candida* cells, the numbers are reduced compared to control. For the control a large cluster of *Candida* cells, sticking to the culture dish surface is seen as shown in figure 2. For the peptide treated *Candida* cells, the numbers got reduced and the cells did not adhere to the surface. These results show that the peptides interfere in biofilm formation. Observing that the AMPs have lytic activity and did not permit the *Candida* cells to adhere on the surface, we postulated that the AMPs may bind to the cell surface glycol-proteins responsible for adhering to the host cells to form biofilm and inhibits the yeast to hyphal transition which is involved in invasion off host mucosal epithelial cell and tissue damage. Hence, we computationally predicted the binding sites of the peptides to Exo-β -(1,3)-Glucanase, Secreted aspartic proteinase (Sap) 1 and N-terminal domain of *Candida albicans* Als9-2. These three proteins are responsible for binding of yeast to host cell surface and meditate infections.

Exo-β-(1,3)-glucanase enzyme is a virulence factor that degrades the *Candida* β-1,3-glucans in the cell wall which helps to invade host tissues and evade the immune system. It also reshapes the *Candida* cell wall for infecting the host cells. Secreted aspartic proteinases (SAPs) are a group of enzymes that play a critical role in the virulence and pathogenicity of *Candida* species. They are responsible for nutrient acquisition and tissue penetration. Agglutinin-Like Sequence 9 (ALS 9) are group of *Candida* adhesin superfamily that play an important role in adhesion, biofilm formation, and immune evasion leading to virulence of host cells.

Computational prediction shows the peptides have strong affinity to *Candida* cell wall proteins which may in turn block the pathogen to bind on the host cells. The three mentioned peptides have unique binding sites that interact with specific amino acids to disrupt the biofilm formation in varying degree.

After determining that the AMPs inhibit the growth of *Candida* cells, the mechanism of disruption was analyzed by scanning electron microscopy (SEM). It shows that the peptide treated *Candida* cells have pores on the membrane. This may lead to the loss of fungal cytoplasmic contents including metabolite and ions by forming deep pits. Further, the loss of turgor and transmembrane potential were the likely causes of cell shrinkage [40, 41]. After confirming that the peptides penetrated through the membrane, their ability to induce Reactive Oxygen Species (ROS) was tested against *C. albicans*. The *C. albicans* was incubated with peptides for 24 h. At this point, the peptides killed the *Candida*, and they did not stick to the plates because their adherence potential was disrupted. However, for the leftover adherent *Candida* cells, there was more ROS generation as seen by increased green fluorescence of DCHFDA which was not seen in the control. The epinecidin-1 and its variants induce ROS generation like other AMPs previously reported from plant defensin [42].

The present study demonstrates about the efficacy of the antimicrobial peptide epinecidin-1 and its variants against *Candida* pathogens bioactivity and biofilm formation. Further characterization is required to find the mode of action at molecular level with and without antifungal agents, which are under pipeline in our laboratory.

## 5. Conclusion

This is the first report on lysine-substituted variants of epinecidin-1 demonstrating anti-*Candida* and antibiofilm effects. The variants showed two-to-eight-fold enhanced activity than the wild type epinecidin-1.

## 6. Abbreviations

AMP: Anti microbial Peptide
CDC: Centre for Disease Control
CFU: Colony Forming Units
DCFHDA: 2′,7′-dichlorofluorescein diacetate
PBS: Phosphate Buffered Saline
ROS: Reactive Oxygen Species
SSA: Stress-Seventy subfamily A

## 7. Funding and Acknowledgements

This research was funded by the Department of Biotechnology, India, (**Ref. No**. BT/PR2071/BBE/117/241/2016). RUSA 2.0 Biological Sciences (**Ref. No**. 311/RUSA(2.0)/2018), Bharathidasan University, Tiruchirappalli and Tamil Nadu State Council for Higher Education (TANSCHE) (**Ref. No**. 01706/P6/2021). S.J acknowledges Indian Council of Medical Research (ICMR-NET-61754/2010) for granting fellowship. S.R thanks DST-PURSE Department of Science and Technology for fellowship. We thank National College Tiruchirappalli, India for performing Scanning Electron microscopy studies.

## Conflicts of Interest

The authors declare no conflict of interest.

**Supplementary Table 1.** Amino acids of the *Candida* surface proteins that interacts with the peptides. The amino acids of the *Candida* membrane proteins that exhibit shared interactions with the peptides are indicated in bold.

|  | *Candida* protein Name | PDB ID | Epi | Var-1 | Var-2 |
| --- | --- | --- | --- | --- | --- |
| a | Exo-B-(1,3)-Glucanase | 1CZ1 | VAL 4<br>TRP 8<br>ARG 16<br>ASN 126<br>ASN 127<br>THR 357 | ASP 151<br>TYR 153<br>VAL 197<br>PHE 229<br>**GLN 230**<br>**PHE 232**<br>GLU 262<br>ASP 280<br>ASN 305<br>ARG 309 | PRO 196<br>**GLN 230**<br>VAL 231<br>**PHE 232**<br>ARG 309<br>TYR 317<br>ASP 318 |
| b | Secreted aspartic proteinase | 2QZW | ARG 54<br>GLY 220<br>**THR 221**<br>THR 222<br>**TYR 225**<br>GLN 228<br>**HIS 249**<br>TYR 252<br>SER 300<br>ILE 301 | LYS 49<br>GLY 85<br>ASP 86<br>GLY 87 | HIS 11<br>VAL 12<br>**THR 221**<br>**TYR 225**<br>GLY 248<br>**HIS 249**<br>GLU 278<br>ALA 282<br>GLN 295<br>LEU 298<br>ARG 312 |
| c | N-terminal domain of Als 9-2 | 2Y7L | TYR 21<br>VAL 22<br>GLY 165<br>TYR 166<br>CYS 188<br>GLU 189<br>CYS 280<br>GLY 283<br>ASN 284<br>ASP 288<br>THR 291 | **THR 63**<br>**THR 65**<br>ASN 82<br>PHE 87<br>PRO 174<br>**SER 175**<br>ASN 177<br>**ILE 267**<br>ASN 299 | **THR 63**<br>**THR 65**<br>ASP 80<br>SER 92<br>**SER 175**<br>GLU 265<br>**ILE 267**<br>ASP 268 |

